# Transcriptome-wide Mendelian randomization study prioritising novel tissue-dependent genes for glioma susceptibility

**DOI:** 10.1101/2020.10.12.335661

**Authors:** Jamie W Robinson, Richard M Martin, Spiridon Tsavachidis, Amy E Howell, Caroline L Relton, Georgina N Armstrong, Melissa Bondy, Jie Zheng, Kathreena M Kurian

## Abstract

Genome-wide association studies (GWAS) have discovered 27 loci associated with glioma risk. Whether these loci are causally implicated in glioma risk, and how risk differs across tissues, has yet to be systematically explored.

We integrated multi-tissue expression quantitative trait loci (eQTLs) and glioma GWAS data using a combined Mendelian randomisation (MR) and colocalisation approach. We investigated how genetically predicted gene expression affects risk across tissue type (brain, estimated effective n=1,194 and whole blood, n=31,684) and glioma subtype (all glioma (7,400 cases, 8,257 controls) glioblastoma (GBM, 3,112 cases) and non-GBM gliomas (2,411 cases)). We also leveraged tissuespecific eQTLs collected from 13 brain tissues (n=114 to 209).

The MR and colocalisation results suggested that genetically predicted increased gene expression of 12 genes were associated with glioma, GBM and/or non-GBM risk, three of which are novel glioma susceptibility genes (*RETREG2/FAM134A, FAM178B* and *MVB12B/FAM125B*). The effect of gene expression appears to be relatively consistent across glioma subtype diagnoses. Examining how risk differed across 13 brain tissues highlighted five candidate tissues (cerebellum, cortex, and the putamen, nucleus accumbens and caudate basal ganglia) and four previously implicated genes (*JAK1, STMN3, PICK1* and *EGFR*).

These analyses identified robust causal evidence for 12 genes and glioma risk, three of which are novel. The correlation of MR estimates in brain and blood are consistently low which suggested that tissue specificity needs to be carefully considered for glioma. Our results have implicated genes yet to be associated with glioma susceptibility and provided insight into putatively causal pathways for glioma risk.

## Introduction

Gliomas are the largest group of intrinsic brain tumours, with age-adjusted incidence rates ranging from 4.67 to 5.73 per 100,000 [1]. Furthermore, malignant gliomas cause significant years of life lost compared with other cancer types – about 20 years of life lost on average – due to late diagnosis and poor treatment outcomes [2], Broadly, gliomas can be subclassified as glioblastoma (GBM, World Health Organisation (WHO) grade IV), which are the most aggressive subtype with a relatively short clinical overall survival, and what is termed non-GBM, lower-grade WHO grade II and III gliomas which have longer survival times. This lower-grade glioma group has not been precisely defined in our dataset but may be presumed to include mostly diffuse astrocytoma, anaplastic astrocytoma, oligodendroglioma and anaplastic oligodendroglioma. To date, there are only two broadly accepted risk factors for glioma. The first, exposure to ionising radiation, accounts for only a small portion of cases [3]. The second includes rare heritable genetic factors. The latest glioma genome-wide association study (GWAS) identified 27 loci that are associated with glioma risk, but it is estimated that we have uncovered only about a third of the risk posed by familial or inheritable factors (27% for GBM and 37% for non-GBM) [4], indicating a large portion of genetic glioma risk is still to be uncovered.

Investigating and understanding how genes are differentially expressed in tumour subtypes has led to a better understanding of gliomagenesis through potentially related mechanisms and pathways and has also improved clinical outcomes for patients due to differential treatments. Previous studies have shown that genes are differentially expressed in glioma dependent on subtype [5–10]. Furthermore, gene expression profiling was proposed as a better method of diagnosis over the previous clinical practice of histological grading because classification based on gene expression seemed to better predict for survival [6, 8, 11]. In 2016, the WHO classification for central nervous system tumours updated their diagnostic rubric to include analysis of the tumour genome [12], It is likely that in the latest WHO guidelines, which at the time of writing are undergoing consultation, will still include genetic diagnostic and prognostic biomarkers to inform glioma classification and outcome [13]. Whilst including genetic factors into the classification criteria has seen measurable benefits for patients, functional studies have been limited and it is not known if certain mutations are merely correlated with gliomagenesis and subtype differentiation or play a causal role in risk. How genetic markers differ by subtype diagnosis is therefore important both to elucidating mechanisms of glioma risk and development, and to further improve clinical outcomes for patients.

In this study, we utilise Mendelian randomisation (MR) – an established instrumental variable method – to assess the causal relationship between genetically predicted gene expression on glioma subtype risk [14], MR suffers less from biases, such as reverse causation and confounding, that invariably limit causal inference in traditional epidemiological studies [14, 15]. Colocalisation is a complementary statistical method that can identify putative causal genetic variants shared by two traits [16]. It provides orthogonal evidence of causality and strengthens MR findings where genetically predicted gene expression affects glioma subtype risk. Integrating MR analyses with expression data from brain tissues provides insight into how tissue-specific gene expression may differentially alter glioma risk across the brain. These data are linked to eQTLs derived from blood to determine how the risk profile for glioma differs between brain tissue and whole blood.

## Methods

### Data

We used summary-level data from different GWAS to compare eQTLs from brain tissue (estimated effective n = 1,194) [17] and from whole blood (n = 31,684) [18]. Our analysis involved a two-sample MR framework, whereby the exposure and outcome data comprise independent populations, to estimate the causal effect of gene expression variation on glioma risk (based on subtype diagnoses of all glioma, GBM and non-GBM). In follow-up sensitivity analyses, we used eQTLs from Genotype-Tissue Expression version 8 (GTEx v8, https://gtexportal.org/home/ [19]) (n = 114 to 209) [20] to examine tissue-specific effects of gene expression. **Table 1** summarises the datasets used in this analysis. The glioma data were based on a meta-analysis of three glioma GWAS consisting of 7,400 glioma cases and 8,257 controls (Glioma International Case-Control Study (GICC), MD Anderson Study (MDA) and GliomaScan datasets [4]). These include 3,112 GBM cases and 2,411 non-GBM cases – the remaining 1,877 glioma cases did not have subtype diagnosis available.

**Table 1.** List of datasets used in this study. The meta-analysis is taken from Qi, *et al*. [17] which includes data from GTEx v6 [20], the Religious Order Study and Memory and Ageing Project (ROSMAP) [32] and CommonMind Consortium (CMC) [33]. eQTLGen Consortium (eQTLGen) [34] contains eQTLs from whole blood. GTEx v8 data is used to interrogate tissue-specific expression [20].

| Dataset | Tissue | No. of Genes | Sample Size |
| --- | --- | --- | --- |
| Meta-analysis | Brain | 8,277 | 1,194 |
| eQTLGen | Whole blood | 19,942 | 31,684 |
| GTEx v8 | Amygdala | 3,726 | 129 |
|  | Anterior cingulate cortex (BA24) | 5,640 | 147 |
|  | Caudate (basal ganglia) | 8,362 | 194 |
|  | Cerebellar hemisphere | 10,027 | 175 |
|  | Cerebellum | 11,240 | 209 |
|  | Cortex | 9,082 | 205 |
|  | Frontal cortex (BA9) | 7,335 | 175 |
|  | Hippocampus | 5,517 | 165 |
|  | Hypothalamus | 5,499 | 170 |
|  | Nucleus accumbens (basal ganglia) | 8,198 | 202 |
|  | Putamen (basal ganglia) | 6,902 | 170 |
|  | Spinal cord (cervical c-1) | 4,483 | 126 |
|  | Substantia nigra | 3,301 | 114 |

### Instrument Selection

We constructed instruments from each dataset **(Table 1**) using independent (*r*^2^ < 0.01) SNPs that met genome-wide significance (P < 5.00×10^-8^). We pre-specified that results must meet a strict Bonferroni-corrected P value threshold of 7.30×10^-6^ (0.05 / 6,849, the number of tests ran) or a suggestive P value threshold of 9.49×10^-5^ (0.05 / 6,849, multiplied by 13 for each tissue type). eQTLs were categorised based on whether they were *cis*-acting or *trans*-acting, defined as SNPs within and without a 1Mbp window of the gene regulatory region, respectively. We included only *cis*-acting eQTLs in our analysis as *trans*-acting eQTLs are more prone to horizontal pleiotropy due to their distal nature on the genome.

### Identifying the Causal Effects of Genetically Predicted Gene Expression on Glioma Risk

For the main MR analysis, we applied two-sample MR to estimate the causal relationship between eQTLs and glioma using the MR-Base R package [21], Most (86%) of tests consisted of the Wald ratio MR method as many eQTLs were instrumented by a single SNP; eQTLs that were instrumented by multiple SNPs were analysed using the inverse variance weighted (IVW) method. We obtained MR results associated with all glioma, GBM and non-GBM risk. Each eQTL that met at least the suggestive P value threshold (P < 9.49×10^-5^) had a region of ±500Kbp around the instrumented SNP(s) extracted, which was subjected to a conditional analysis using Conditional and Joint analysis (GCTA-COJO) [22] to ensure only independent signals within the region remained before colocalisation using the coloc R package [16]. The coloc package provides estimates for five hypotheses related to whether a single causal variant is shared between two traits. The final hypothesis, H_4_, indicates the percentage chance the tested variant is causal and shared between both traits. Throughout we provide the colocalisation estimates in regard to the H_4_ hypothesis as this is the only estimate required for our analyses. Further details about the coloc R package are given by Giambartolomei, *et al*. [16].

Finally, we also applied the Steiger filtering method to ensure results were not distorted due to the presence of reverse causation [23]. Results from the Steiger filtering analysis are presented as a categorical variable to aid comprehension: true, if the direction of effect is from exposure to outcome and P < 0.05; false, if the direction of effect is reversed and P < 0.05; uncertain if P ≥ 0.05.

### Examining Tissue-Specific Effects of Gene Expression on Glioma Risk

We included each MR association which passed the suggestive P value threshold (P < 9.49×10^-5^) in a follow-up sensitivity analysis to examine tissue-specific effects of gene expression on all glioma, GBM and non-GBM risk. Genes were systematically mapped to relative genes across 13 brain tissues from GTEx v8 based on Ensembl IDs (ENSG). We selected instruments from independent (r^2^ < 0.01) SNPs that met a lenient threshold (P < 5.00×10^-4^) to ensure a greater chance that there will be an eQ,TL for each tissue type. This threshold should be viewed as enabling a heuristic approach to the tissue-specific analyses by allowing for more genes to be instrumented in different tissues [24], Only c/s-acting eQTLs were included to avoid potentially pleiotropic *trans*-acting eQTLs. We analysed these data using the MR-Base R package and compared the magnitude and direction of the effect estimate across tissue types and subtype diagnosis.

To ascertain whether tissue-specific gene expression differentially altered risk of glioma, GBM and non-GBM, we conducted heterogeneity analyses using Cochran’s Q test. We also calculated the tau-score, a quantitative measure of tissue-specificity derived by Kryuchkova-Mostacci and Robinson-Rechavi [25]. This method can be naively summarised as summing the weighting of a gene’s expression in a single tissue against the maximum expression over all tissues. The tau-score will range between 0 and 1, where 0 means the gene is broadly expressed and 1 means specific expression; in their paper, Kryuchkova-Mostacci and Robinson-Rechavi define a threshold cut-off for specific expression at 0.8 which we also use [25]. Finally, we constructed Z-scores for each SNP to determine which SNPs in which tissues may be driving heterogeneity in the results.

We hypothesised that due to the presence of the blood-brain barrier, brain-based eQTLs and blood-based eQTLs should have little correlation with one another. We systematically linked causal estimates for eQTLs that appeared in both brain tissue and blood. We then compare these data to determine whether blood-based eQTLs were correlated with brain-based eQTLs and whether easier-to-gather blood data could proxy sufficiently for brain data.

## Results

In total, our MR analysis of brain and blood eQTLs identified 34 associations that met at least the suggestive P value threshold (P < 9.49×10^5^) for 17 genes associated with risk of glioma, GBM or non-GBM **(Figure 1**). Altogether, six genes were instrumented by eQTLs in blood and 12 genes had eQTLs from brain tissue -one gene, *JAK1*, had an associated eQTL in both brain and blood. We found that 20 associations had strong evidence (H_4_ ≥ 80%, see **Methods** for explanation) of colocalisation. Steiger filtering revealed the direction of the causal estimate was correctly orientated from gene expression to subtype diagnosis in 29 associations; the remaining five showed an uncertain result due to the P value for Steiger filtering not reaching 0.05. Overall, 17 tissue-subtype associations for 12 genes showed robust causal evidence from the MR and colocalisation analyses and passed the Steiger filtering analysis. These 17 associations and 12 genes formed our main results and were subjected to the tissue-specific analyses. Results are presented in **Table 2** (with further details in **Supplementary Table 1**).

**Figure 1.**
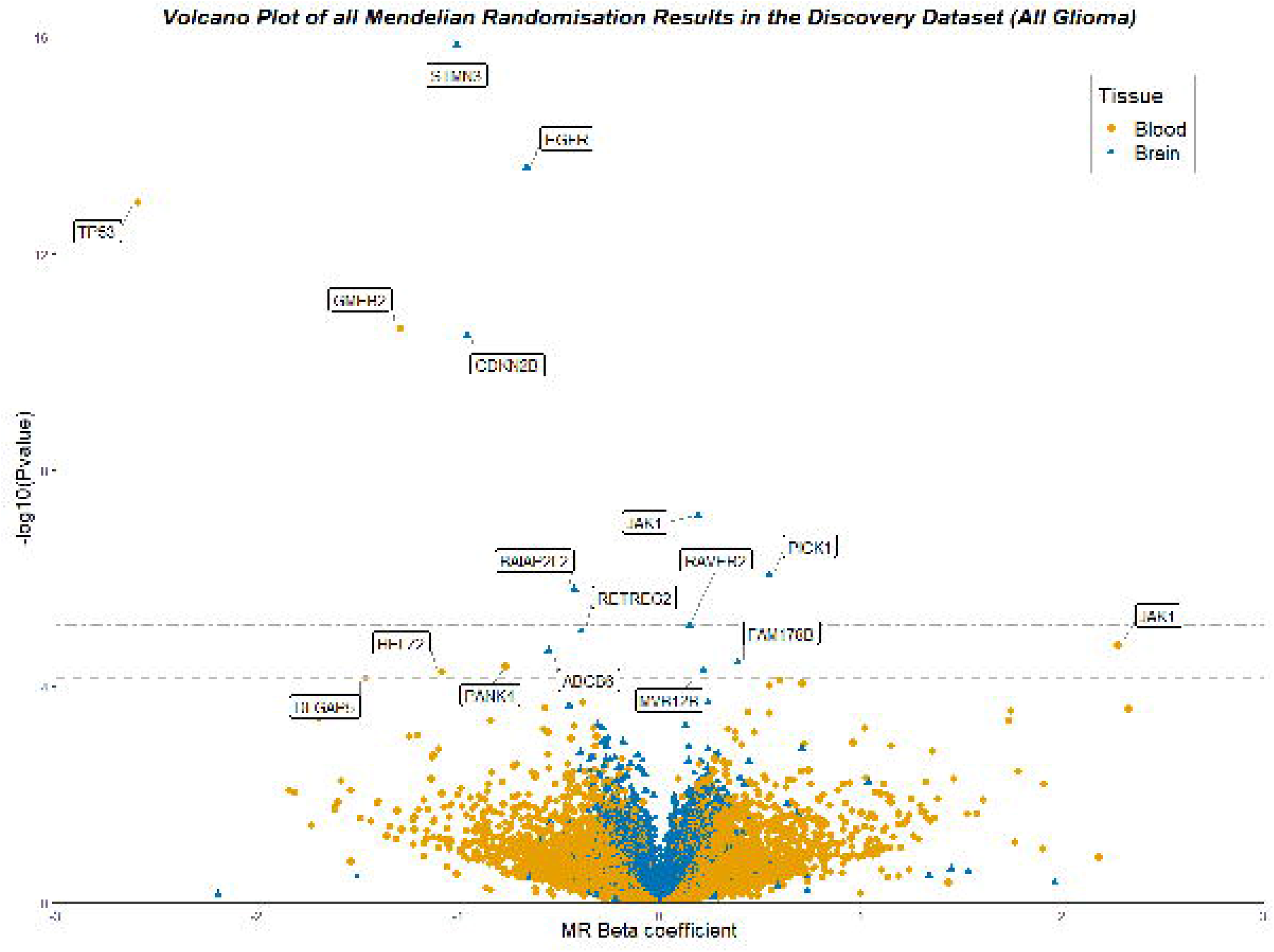
Volcano plot of all results from the main MR analysis of brain and blood eQTLs and all glioma. The horizontal dot-dashed line represents the Bonferroni-corrected P value threshold (P < 7.30×10^-6^) and the horizontal dashed line is the suggestive P value threshold (P < 9.49×10^-5^). Genes are labelled if they pass at least the suggestive threshold.

**Table 2.** Main associations which showed robust evidence from the Mendelian randomisation, colocalisation and Steiger filtering analyses. The colocalisation result for a single, shared causal variant between the gene and all glioma is provided (“Coloc (%)”, see **Methods).** Steiger filtering showed the correct orientation for the direction of effect between gene expression and subtype risk for all results in this table.

| Gene | SNP(s) | Tissue | Subtype | OR (95% CI) | P value | Coloc (%) | Steiger P value |
| --- | --- | --- | --- | --- | --- | --- | --- |
| **ABCB6** | rs75450661 | Brain | All glioma | 0.57 (0.44, 0.74) | $2.20 \times 10^{-5}$ | 97 | $7.41 \times 10^{-6}$ |
| **BAIAP2L2** | rs1004764 | Brain | All glioma | 0.65 (0.55, 0.78) | $1.62 \times 10^{-6}$ | 96 | $2.36 \times 10^{-10}$ |
| | | | GBM | 0.60 (0.49, 0.73) | $2.85 \times 10^{-7}$ | 81 | $1.25 \times 10^{-9}$ |
| **EGFR** | rs6979446,<br>rs759170 | Brain | GBM | 0.45 (0.38, 0.53) | $9.99 \times 10^{-20}$ | 81 | $3.53 \times 10^{-6}$ |
| **FAM178B** | rs13407036 | Brain | All glioma | 1.47 (1.23, 1.77) | $3.59 \times 10^{-5}$ | 94 | $1.97 \times 10^{-16}$ |
| **JAK1** | rs2780902 | Brain | All glioma | 1.21 (1.13, 1.29) | $6.95 \times 10^{-8}$ | 81 | $6.89 \times 10^{-139}$ |
| | | | GBM | 1.27 (1.17, 1.37) | $1.56 \times 10^{-9}$ | 95 | $1.84 \times 10^{-134}$ |
| **MVB12B** | rs4837096 | Brain | All glioma | 1.24 (1.12, 1.38) | $5.27 \times 10^{-5}$ | 97 | $2.53 \times 10^{-23}$ |
| **PANK4** | rs2985862 | Blood | All glioma | 0.46 (0.32, 0.67) | $4.30 \times 10^{-5}$ | 97 | $4.62 \times 10^{-10}$ |
| **PICK1** | rs5756894 | Brain | All glioma | 1.72 (1.39, 2.14) | $8.82 \times 10^{-7}$ | 97 | $4.34 \times 10^{-7}$ |
| | | | GBM | 1.96 (1.54, 2.51) | $6.60 \times 10^{-8}$ | 92 | $2.13 \times 10^{-6}$ |
| **PRLR** | rs67975005 | Brain | All glioma | 0.66 (0.54, 0.82) | $9.33 \times 10^{-5}$ | 91 | $1.13 \times 10^{-7}$ |
| **RETREG2** | rs1996719 | Brain | All glioma | 0.68 (0.57, 0.80) | $9.54 \times 10^{-6}$ | 98 | $7.90 \times 10^{-11}$ |
| | | | GBM | 0.67 (0.55, 0.81) | $6.13 \times 10^{-5}$ | 95 | $9.91 \times 10^{-11}$ |
| **STMN3** | rs6011016 | Brain | All glioma | 0.36 (0.29, 0.46) | $1.44 \times 10^{-16}$ | 96 | $1.44 \times 10^{-16}$ |
| | | | GBM | 0.29 (0.22, 0.38) | $4.55 \times 10^{-19}$ | 97 | $7.88 \times 10^{-3}$ |
| **TP53** | rs35850753 | Blood | Non-GBM | 0.17 (0.09, 0.32) | $9.61 \times 10^{-8}$ | 98 | $3.35 \times 10^{-2}$ |

Comparing our results with previously identified GWAS associations (noted in a review conducted by Kinnersley, *et al*. [26]) revealed that *RETREG2 (FAM134A), FAM178B* and *MVB12B* (*FAM125B*) are putative novel genes implicated in glioma risk that are not also located on a known glioma risk locus and formed part of our main results. The remaining results have been previously implicated in glioma risk through GWAS associations or are located on a known susceptibility locus.

**Figure 2** shows the MR effect estimates for each of the 12 genes and all glioma, GBM and non-GBM subtypes. The direction and magnitude of the estimated causal effect broadly agreed across all genes and subtypes. However, the non-GBM results were noticeably attenuated, for example, in the case of *JAK1*.

**Figure 2.**
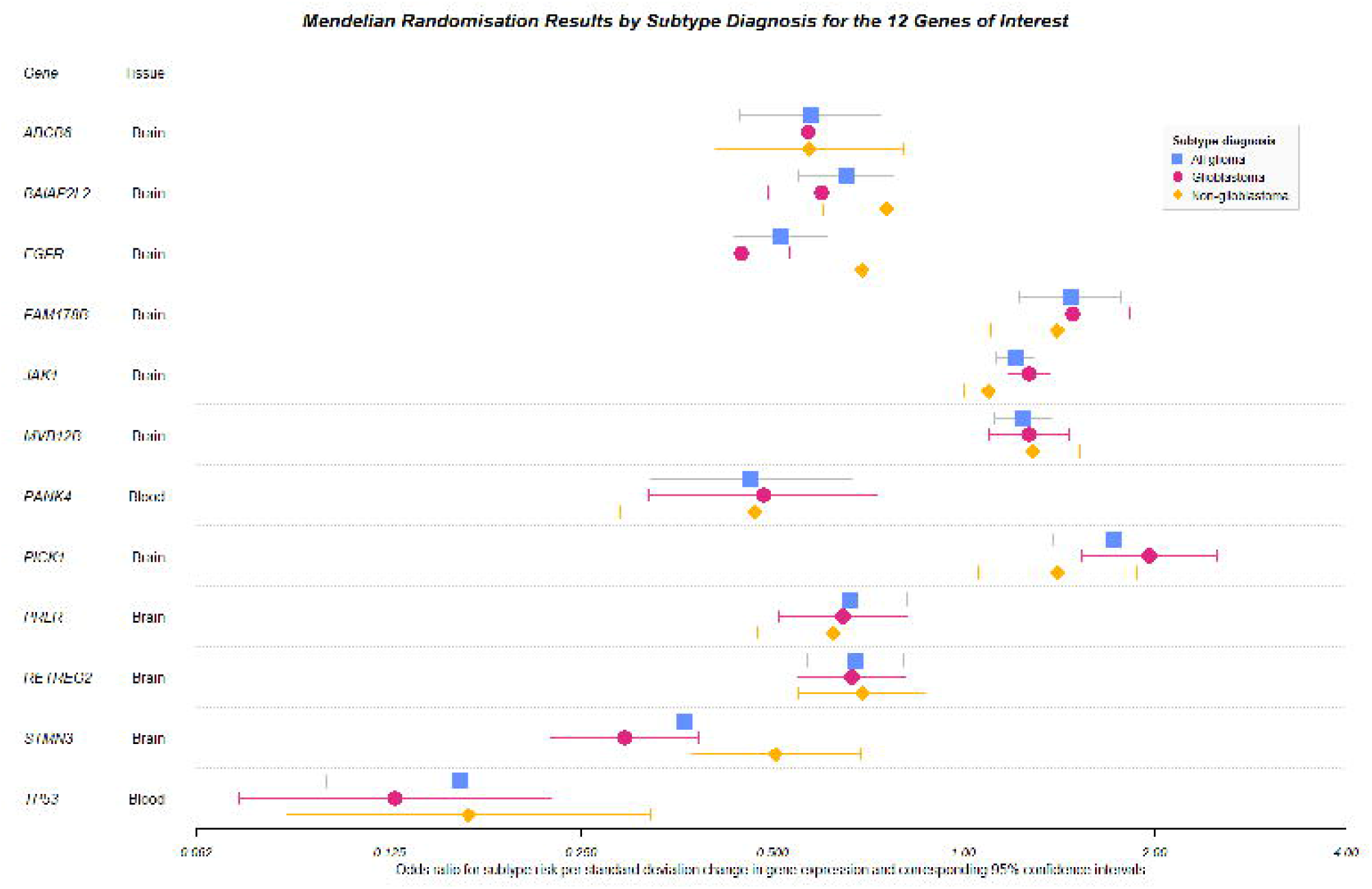
Forest plot of Mendelian randomisation results for the 12 genes which had robust MR and colocalisation evidence and also passed the Steiger filtering analysis.

To examine how tissue-specific gene expression affected glioma risk, the 12 genes which formed our main results were systematically linked to eQTLs across 13 brain tissues using GTEx v8. The effects of tissue-specific expression of most genes were assessed using MR in 8 to 13 tissues (mean = 10 tissues) except for *ABCB6,* which had data in four tissues **(Supplementary Table 2).** Full results for the tissue-specific MR analysis are in **Supplementary Table 3a.** These results broadly agreed with the main MR analysis, though results were attenuated due to the smaller sample sizes present in the GTEx v8 dataset. Applying the same threshold for the discovery MR analysis (P < 7.30×10^-6^) revealed that 56% of the results arise in five tissues: putamen (basal ganglia) (12%), cortex (11%), cerebellum (11%), caudate (basal ganglia) (11%) and nucleus accumbens (basal ganglia) (11%) **(Supplementary Table 3b).** Furthermore, 100% of these results arose due to four genes: *STMN3* (32%), *EGFR* (26%), *PICK1* (23%) and *JAK1* (18%) **(Supplementary Table 3c).**

Two tissue-specific results showed evidence of high heterogeneity. These were *EGFR* for all glioma (Cochran’s Q= 155.96, P = 1.72×10^-28^) and GBM (Cochran’s Q= 162.38, P = 3.49×10^-27^) subtype analyses **(Supplementary Table 4).** Examination of the *EGFR* SNPs’ Z-scores highlighted three tissues (hippocampus, hypothalamus and substantia nigra) whereby the effect of the instrumented SNP was in the opposite direction (positive Z-scores) compared to the remaining 10 SNPs in other tissues (negative Z-scores) **(Supplementary Table** 5). Only *PICK1* showed potential tissue-specific expression with a tau-score of 0.78 **(Supplementary Table 4).**

Finally, we compared effect estimates between of the estimated effects from the MR analysis of the same gene expressed in brain and blood tissues on glioma risk. We applied four P value thresholds (P < 0.1, 0.05, 0.01, 0.005) to examine whether the strength of the MR association influences the correlation between estimates in brain and blood tissues. Overall, we observed a low correlation between brain and blood eQTLs (Pearson correlation = 0.18, number of genes = 632) at the highest P value threshold (P < 0.1). After applying the more stringent threshold (P < 0.005), the correlation increased but remained low (Pearson correlation = 0.21, number of genes = 45). These results are shown in **Figure 3.**

**Figure 3.**
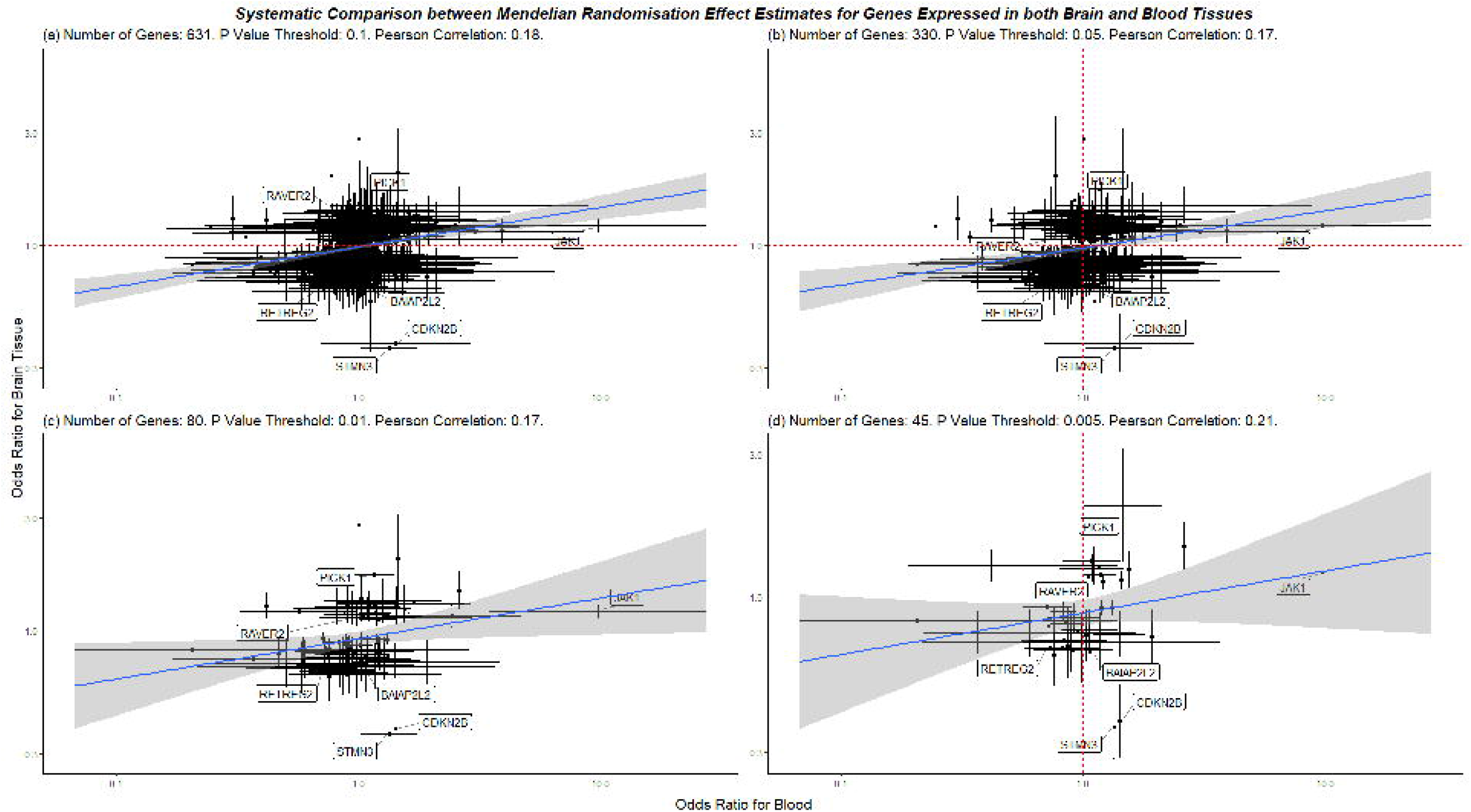
Systematic comparison between the MR results from brain tissues and blood. Any eQTL that appeared in both brain and blood datasets was included in this analysis. We plotted the odds ratios for blood against brain, deciding which results to include based on a P value cut-off: **(a)** P < 0.1, **(b)** P < 0.05, **(c)** P < 0.001 and **(d)** P < 0.005. Labels are provided for genes which had an association that passed at least the suggestive P value threshold (P < 9.49×10^-5^).

## Discussion

In this study, we combined MR and colocalisation analyses to estimate the genetically predicted gene expression on glioma risk which provided causal evidence for 12 genes. Three of these genes are novel in the context of glioma risk. Overall, these results were robust to sensitivity analyses, including Steiger filtering and tissue-specific analyses.

*RETREG2* (or *FAM134A*), *FAM178B* and *MVB12B* appear to be novel findings related to glioma risk. Reticulophagy Regulator Family Member 2 (*RETREG2, FAM134A*) is a protein-coding gene whose function is largely unknown. Examination of the Human Protein Atlas reveals that expression of *RETREG2* RNA and protein is primarily located in the brain and testes, though tissue specificity is low [27]. Family with Sequence Similarity 178 Member B (*FAM178B*) is another protein-coding gene with an undocumented function. It is an important paralogue of gene *SLF2* whose protein plays a role in the DNA damage response. Finally, Multivesicular Body Subunit 12B (*MVB12B, FAM125B*) is a regulator of vesicle trafficking and has been implicated in lipid and ubiquitin binding. Overexpression of this gene and its protein inhibits HIV-1 infectivity by regulating ESCRT (endosomal sorting complex required for transport-I)-mediated virus budding [28]. A 2014 study created a nine-gene-signature panel, which included *MVB12B,* that accurately predicted prognosis for glioma patients, further implicating the gene’s role in glioma biology [29]. Further research into these genes is warranted to provide replication and to elucidate potential pathways by which these genes affect glioma risk.

When considering differential risk across subtype diagnosis, our MR results showed agreement in the direction of effect for risk of all glioma, GBM and non-GBM. However, we also found that associations with non-GBM risk tended to be weaker in magnitude than associations with the other two subtypes of glioma and are generally attenuated. Case numbers are broadly similar – 3,112 GBM cases versus 2,411 non-GBM cases - but may still be underpowered in the non-GBM analysis. This is evidenced by the consistently larger P values for the MR results of the non-GBM analysis when compared to the GBM analysis. Whether this attenuation is due to the heterogeneous nature of brain tumours, or due to the lack of power in the subtype analysis, requires follow-up analyses in larger datasets. Overall, however, there is little evidence to conclude there exists a large difference in the gene expression risk profile of GBM and non-GBM tumours. Therefore, future studies in this area may seek to consolidate data independent of subtype diagnosis such that larger statistical power may be achieved.

Gliomas may develop across the entirety of the central nervous system but are generally found in the cerebrum, particularly the frontal and temporal lobe, and less commonly in the cerebellum depending on the age of the patient [30]. We sought to determine how genetically predicted gene expression in the 13 brain tissue types collected by GTEx v8 [19] compared to the anatomical regions within which tumours are found. The 12 genes that formed our main results were matched to, on average, an eQTL in 10 brain tissues allowing for a broad investigation on how glioma risk is affected by gene expression in disparate tissues. Applying MR and a similar P value threshold (P < 7.30×10^-6^) revealed that 56% of the results that met that threshold arise in five of the 13 tissues. Two of these are common/uncommon tissues for glioma (cortex (11%), cerebellum (11%), respectively). The other three tissues were from the deep brain and are considered rarer locations for glioma (putamen (basal ganglia) (12%), caudate (basal ganglia) (11%) and nucleus accumbens (basal ganglia) (11%)) **(Supplementary Table 3b)**. These analyses provided evidence that gene expression in these five tissues potentially drives glioma risk and, owing to the diffusive nature of the tumours, are then found elsewhere in the brain. Furthermore, 100% of these results arose due to four genes: *STMN3* (32%), *EGFR* (26%), *PICK1* (23%) and *JAK1* (18%) **(Supplementary Table 3c)** highlighting these genes of high importance for follow-up studies. *EGFR* also showed high heterogeneity for all glioma and GBM subtypes risk, indicating gene expression in certain tissues may affect risk differently – for *EGFR* these were the hippocampus, hypothalamus and substantia nigra. However, our analyses showed little evidence of tissue-specific gene expression, with one gene showing suggestive evidence of tissue specificity (*PICK1*, tau-score = 0.78). Larger sample sizes will allow us to clarify exactly how gene expression across multiple tissues differentially affects glioma risk. Broadly we have shown in our analyses that gene expression across the entire brain, agnostic of tissue site, drives glioma risk as opposed to gene expression specific to certain tissues, though the same gene expressed in different tissues may differentially affect risk.

We also investigated whether blood eQTLs, for which there are datasets of large sample sizes available, can proxy for relatively low powered brain tissue eQTLs. We compared how the MR effect estimates differed between eQTLs that were systematically linked between datasets. We found that even after applying an increasingly stringent P value threshold, there is little correlation between the MR associations for brain and blood in the context of glioma risk **(Figure 2).** We interpreted these results to mean that should gene expression in brain be associated with glioma risk, the same gene expression in blood cannot be assumed to also affect risk similarly; in some cases, the risk profile of genes expressed in brain and blood differed in direction of effect, e.g. *STMN3* seems to increase risk when expressed in blood and decrease risk when expressed in brain. A potential avenue of future research is to determine why some genes, like *STMN3,* differentially affect risk depending if they are expressed in brain or blood.

Strengths of our analysis include the use of genetic variants that proxy for gene expression levels, which should reduce the influence of confounding and bias through reverse causation. Furthermore, these genetic variants were obtained from relatively large meta-analyses allowing for increased statistical power and more precise estimates. The addition of subtype diagnoses and tissue-specific data has allowed us to deeper investigate the risk profile of glioma.

Our methodology is also a strength of our analysis. MR is less liable to sources of confounding and bias, and provides evidence of causal relationships between genetic expression and glioma risk. Combining the MR results with colocalisation provides further evidence of causality by determining whether gene expression and glioma risk share a single, causal variant or through distinct causal variants that are in linkage disequilibrium with one another. Follow-up analyses in different tissues also act as a replication study providing further evidence of causality. However, despite evidence of causality, this study does not prove causality and further studies would be required to determine this.

Our study is not without limitations. Despite the use of relatively large datasets for the traits we studied, i.e. gene expression and glioma, our analyses still likely suffer from low statistical power arising from small sizes particularly evidenced in the non-GBM subtype analyses. All but one of our main results were instrumented by a single SNP which limited our ability to undertake common MR sensitivity analyses to detect, for example, horizontal pleiotropy, a common source of confounding in MR studies. Colocalisation has been proffered as a sensitivity analysis that can at least eliminate spurious associations that have arisen due to horizontal pleiotropy because a shared causal variant for two traits is necessary, though not sufficient, for them to be causally related [23]. Despite this, horizontal pleiotropy remains a concern for QTL studies like ours due to instruments generally consisting of single SNPs. Another limitation to our study is that MR will provide effect estimates for lifetime exposure to gene expression whereas expression levels of genes can frequently change and even within glioma, cells at the leading edge of the tumour appear to exhibit a different expression profile to those at the core [31]. Overall, whilst our results are consistent across the sensitivity analyses we performed, there remains a possibility that they were biased through confounding and horizontal pleiotropy.

We demonstrated the effectiveness of MR and colocalisation to identify putatively causal genes for glioma susceptibility. Our study has revealed causal evidence for three novel genes (*RETREG2, FAM178B* and *MVB12B*) associated with glioma risk. We show that there is no distinct difference between the causal gene expression profile and glioma subtype risk. Finally, our tissue analyses suggested that the causal estimates for glioma are different based on whether the gene is expressed in brain or blood tissue. Finally, our tissue analyses highlight five candidate tissues (cerebellum, cortex, and the putamen, caudate and nucleus accumbens basal ganglia) and four genes (*JAK1, STMN3, PICK1* and *EGFR*) which had causal evidence for affecting glioma risk in further research.

## Declarations

### Competing Interests

The authors declare no competing interests.

## Acknowledgements

We would like to thank GICC, GliomaScan and MDA for access to their data.

## Funding

This work was supported by the Medical Research Council (MRC)/University of Bristol Integrative Epidemiology Unit (IEU) and is supported by the MRC and the University of Bristol (MC_UU_12013/1, MC_UU_12013/2). JWR is supported by a joint studentship from NHS North Bristol Trust and Bristol Tumour Bank (SOCS/SJ1447). JZ is funded by the Vice-Chancellor’s fellowship. CLR and RMM is supported by a Cancer Research UK Programme Grant, the Integrative Cancer Epidemiology Programme (C18281/A19169). JWR, RMM, AEH, CLR and JZ are members of the MRC IEU which is supported by the Medical Research Council and the University of Bristol (MC_UU_12013/1-9). RMM is supported by the National Institute for Health Research (NIHR) Bristol Biomedical Research Centre which is funded by the National Institute for Health Research and is a partnership between University Hospitals Bristol NHS Trust and the University of Bristol. The views expressed in this publication are those of the author(s) and not necessarily those of the NIHR or the UK Department of Health and Social Care.

The funders took no active role in the production of this research.

## Data Availability

The glioma data may be accessed under the European Genome-phenome Archive accession number EGAD00010001657 (https://www.ebi.ac.uk/ega/datasets/EGAD00010001657).

## Notes

### Competing Interest Statement

The authors have declared no competing interest.

